# The role of high cholesterol in age-related COVID19 lethality

**DOI:** 10.1101/2020.05.09.086249

**Authors:** Hao Wang, Zixuan Yuan, Mahmud Arif Pavel, Sonia Mediouni Jablonski, Joseph Jablonski, Robert Hobson, Susana Valente, Chakravarthy B. Reddy, Scott B. Hansen

## Abstract

Coronavirus disease 2019 (COVID19) is a respiratory infection caused by severe acute respiratory syndrome coronavirus 2 (SARS-CoV-2) originating in Wuhan, China in 2019. The disease is notably severe in elderly and those with underlying chronic conditions. A molecular mechanism that explains why the elderly are vulnerable and why children are resistant is largely unknown. Here we show loading cells with cholesterol from blood serum using the cholesterol transport protein apolipoprotein E (apoE) enhances the entry of pseudotyped SARS-CoV-2 and the infectivity of the virion. Super resolution imaging of the SARS-CoV-2 entry point with high cholesterol shows almost twice the total number of endocytic entry points. Cholesterol concomitantly traffics angiotensinogen converting enzyme (ACE2) to the endocytic entry site where SARS-CoV-2 presumably docks to efficiently exploit entry into the cell. Furthermore, in cells producing virus, cholesterol optimally positions furin for priming SARS-CoV-2, producing a more infectious virion with improved binding to the ACE2 receptor. In vivo, age and high fat diet induces cholesterol loading by up to 40% and trafficking of ACE2 to endocytic entry sites in lung tissue from mice. We propose a component of COVID19 severity based on tissue cholesterol level and the sensitivity of ACE2 and furin to cholesterol. Molecules that reduce cholesterol or disrupt ACE2 localization with viral entry points or furin localization in the producer cells, may reduce the severity of COVID19 in obese patients.

## INTRODUCTION

In early 2020, COVID19 spread rapidly throughout the developed world leading to extensive death and an ongoing pandemic. The lethality of SARS-CoV-2 is notably selective for elderly^1,2^ and those with chronic disease such as hypertension, diabetes, Alzheimer’s, cardiovascular disease, and smoking^3–5^ although in the case of smoking not without controvery^6^. Interestingly, almost all children present with very minor symptoms, while elderly and those with underlying conditions experience very severe life-threatening symptoms leading to much higher death rates^7^. This disparity was also observed with SARS-CoV, a closely related virus which failed to kill any children under the age of 12, despite being much more lethal than SARS-CoV-2 in adults^8^. Understanding why young people are resistant in this class of viruses could help both healthy and chronically ill adults to avoid severe symptoms of COVID19.

At least two factors affect SARS-CoV-2 infectivity, 1) the opportunity of the virus to bind to the cell surface and 2) the ability of the virus to enter the cell. SARS-CoV-2 attaches to the cell surface by binding to a receptor, angiotensinogen converting enzyme 2 (ACE2)^9,10^. Once bound, the virus can enter through an apparent cell surface mechanism^10,11^ or through clathrin-mediated endocytosis^12^, a cholesterol dependent mechanism that involves monosialotetrahexosylganglioside1 (GM1) containing lipid clusters (also known as lipid rafts, see Supplemental Figure S1A)^13–15^. Regardless of the pathway, the virus must be primed by proteases^16^. In SARS-CoV-2 there is an efficient furin cleavage site between the S1 and S2 segments that is not found in SARS-CoV^17,18^.

Cholesterol is an insoluble eukaryotic lipid found in membranes throughout the human body, most notably in the plasma membrane^19^. It appears to accumulate in tissue with age^20–22^. Furthermore, chronic inflammation causes cholesterol loading into macrophage rich tissue^23^. Pneumocytes and macrophages in the lung uptake and efflux cholesterol and this process is associated with lung disease^24^ and lung function^25,26^. Recently cholesterol has been shown to be a key factors in SARS-CoV-2 infections^27–30^, but a molecular mechanism for cholesterols effects is unclear. For further discussion on cholesterol in peripheral tissue see Figure S1B.

We recently showed a cholesterol-dependent mechanism for anesthetics that regulates the translocation of membrane proteins between GM1 and PIP2 lipid clusters^31,32^. Previous experiments showed ACE2 can associate with detergent resistant membranes (DRMs)^15,33^ which are similar in composition to GM1 clusters. Here we show ACE2 protein levels are low under native conditions in human lung and levels are low in most cells types including lung. Furthermore, ACE2 translocates between GM1 and PIP2 clusters and increased cholesterol contributes to infectivity and ACE2 localization to GM1 lipids in mouse lung providing a molecular mechanism for cholesterol to regulate SARS-CoV-2 entry.

## RESULTS

### Measurements of cholesterol and ACE2 protein levels in human lung tissue

The level of ACE2 expression and cholesterol have been cited as a reason for SARS-CoV-2 infectivity, but little is known about the relative levels of ACE2 protein and cholesterol in cells types that are easily infected vs not easily infected with the virus. To better compare cells of known infectivity with those of low infectivity we examined cholesterol levels in VeroE6 cells and compared them to HEK293T cells. Vero E6 cells are a kidney cell line that are highly infected with SARS-CoV-2 virus^34^. We also compared cholesterol of smoker’s to cultured carcinoma lung cells A549 and H1793. Human lung cells are susceptible to infection while cultures lungs cells are relatively resistant. We found that cultured cells have between 50-70% less free cholesterol (FC) compared to lung tissue from smokers (Figure 1A).

**Figure 1.**
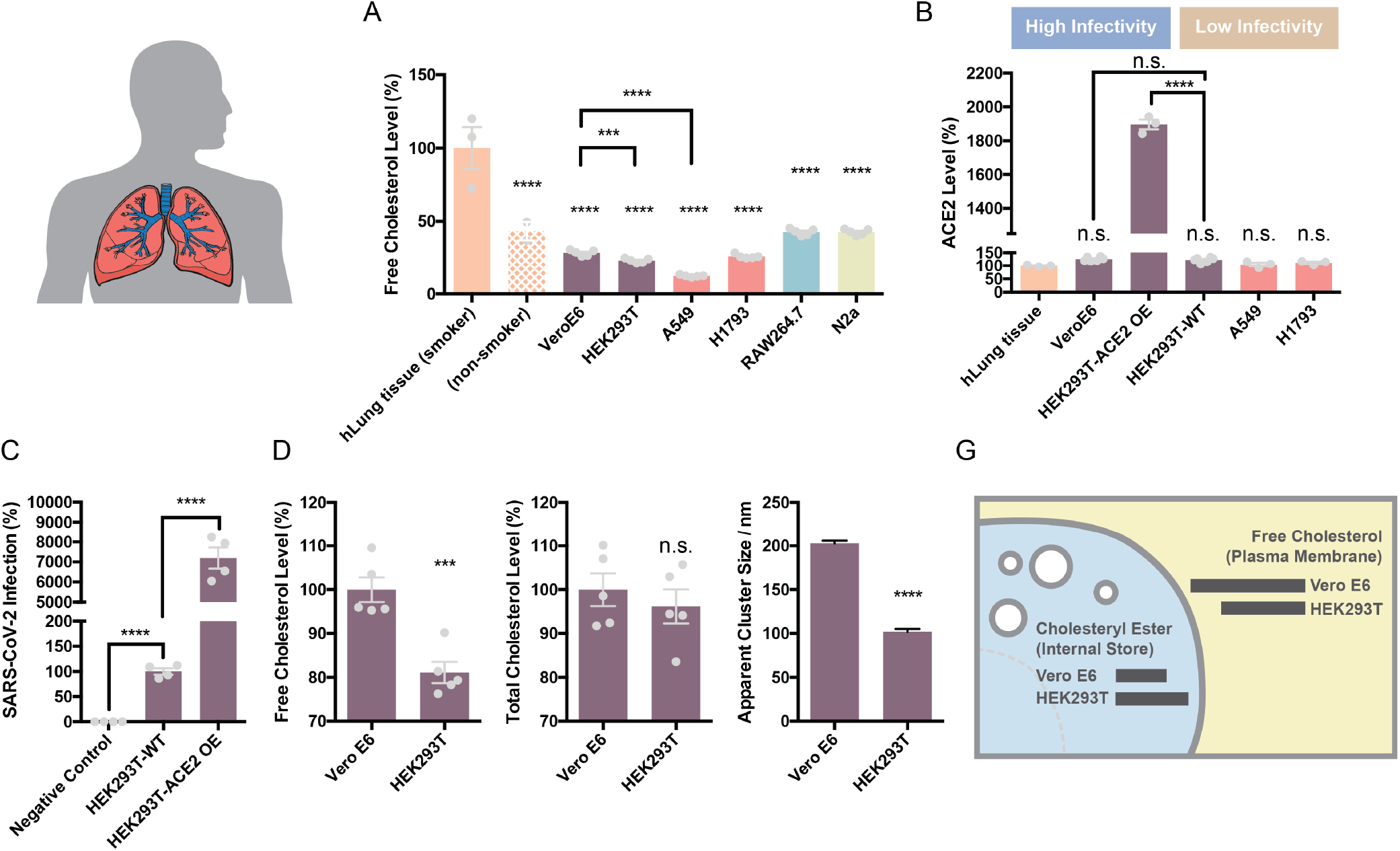
Comparison of free cholesterol (membrane) and ACE2 protein levels. (**A**) Free cholesterol from human lung tissue (epithelial airway; orange, from 3 smokers and 2 non-smokers, including a bronchitis patient and a healthy control) is compared to cultured cells, including kidney cell lines (purple) Vero E6 and HEK293T, lung cell lines (red) A549 and H1793, macrophage cell line RAW264.7 (cyan) and neuroblastoma 2a (N2a) cells. (**B**) ACE2 protein levels in human lung tissue, lung cells and kidney cells measured by ELISA. (**C**) SARS2 PV viral entry with wild type HEK293T expressing endogenous ACE2 (HEK293T WT) and HEK293T cells overexpressing hACE2 (ACE2 OE). Negative (neg.) control are HEK293T cells treated identical to the other conditions except no virus was applied. The endogenous expression was low but presumably much more physiologically relevant and shows ACE2 is expressed in HEK293T sells albeit at comparatively low quantities. The result is a single side by side comparison, but the values are in a typical range we see for these two systems. Data are expressed as mean ± s.e.m., ***P<0.001, ****P<0.0001, one-way ANOVA. (**D-E**) Comparison of free (D) and total (E) cholesterol in cultured Vero E6 and HEK293T cells. Free cholesterol is in the plasma membrane and cholesteryl ester is in the internal store. The free cholesterol is enlarged from panel (A) for clarity. (**F**) Comparison of GM1 lipid clusters size in Vero E6 vs HEDK293 cells using super resolution imaging. GM1 clusters were labeled with fluorescent cholera toxin B. Data are expressed as mean ± s.e.m., ***P<0.001, ****P<0.0001, two-sided Student’s t-test. (**G**) Graphical depiction of the cholesterol levels observed in HEK293T and VeroE6 cells.

**Figure 1.**
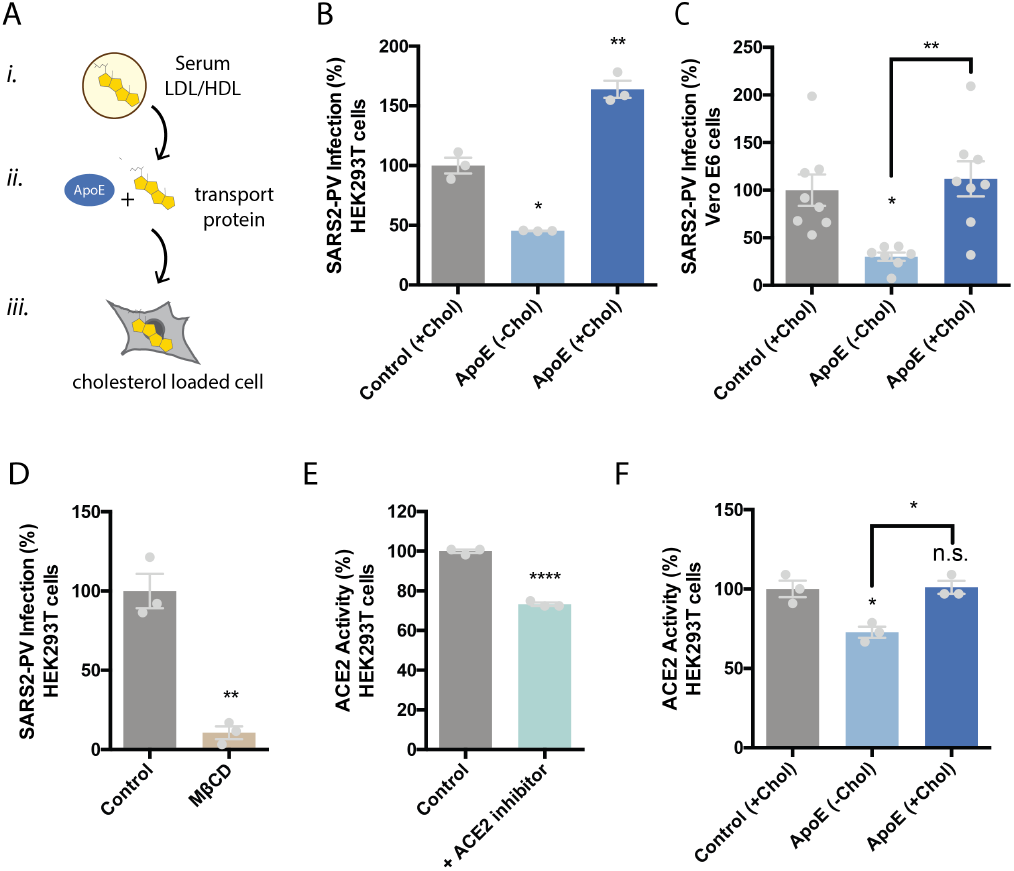
Cholesterol dependent inhibition of SARS-CoV-2 pseudovirus (SARS2-PV) infection. (**A**) Cartoon diagram showing the experimental setup for loading cultured cells with cholesterol. ***i*.,** Cholesterol (yellow shading) loaded into lipoprotein (e.g., low- and high-density lipoprotein (LDL and HDL respectively)) from blood serum. ***ii*.,** Cholesterol free human apolipoprotein E (apoE, brown shading) a cholesterol transport protein is exposed to cholesterol from blood serum and ***iii*,** ApoE transports cholesterol into cells (grey shading) (see also Supplemental Figure S1B). (**B-C**) SARS-CoV-2 pseudovirus (SARS2-PV) entry assay in HEK293T cells (B) and Vero E6 cells (C). Cells were treated with a luciferase expressing retrovirus pseudotyped with the SARS-CoV-2 spike protein that recapitulates viral entry. Infectivity was monitored by a luciferase activity in cells treated with or without apoE. Viral infection in cells with high cholesterol (apoE + serum) was more than 3-fold higher compared to low cholesterol. Data are expressed as mean ± s.e.m., *P<0.05, **P<0.01, one-way ANOVA. (**D**) Depletion of cellular cholesterol with methyl-beta-cyclodextrin (MβCD) blocked almost all viral entry measured by pseudotyped luciferase assay. Data are expressed as mean ± s.e.m., **P<0.01, two-sided Student’s t-test. (**E**) ACE2 activity assay was performed to detect ACE2’s expression in wild type HEK293T cells. Cells incubated with ACE2 inhibitor showed significantly decreased ACE2 activity, suggesting ACE2 is expressed in the HEK293T cells. Data are expressed as mean ± s.e.m., ****P<0.0001, two-sided Student’s t-test. (**F**) ACE2’s enzymatic activity is regulated by membrane cholesterol. ApoE mediated cholesterol depletion from cell membrane inhibits ACE2’s activity. Data are expressed as mean ± s.e.m., *P<0.05, one-way ANOVA.

To test the potential role of increased ACE2 expression in SARS-CoV-2 infectivity, we compared ACE2 protein levels in lung tissue of smokers (epithelial airway) with the various cultured cell types using an ELISA assay for ACE2 protein. We found the highest endogenous levels of ACE2 protein in HEK293T and the lowest in the lung of smokers (Figure 1B) although none were significantly different and all of them were relatively low levels. This is consistent with low levels of endogenous ACE2 in HEK293T cells observed by westernblot^35^, but opposite to those who have used RNA as an indirect surrogate for ACE2 protein expression^36^.

To confirm the presence of endogenously expressed ACE2 in the membrane of WT HEK293T cells, we performed an ACE2 activity assay using a fluorescent substrate cleavage assay. The ACE2 specific inhibitor DX600 decreased fluorescence 70% compared to control (Figure 2E), confirming ACE2 enzyme is present and active in HEK293T cells.

**Figure 2.**
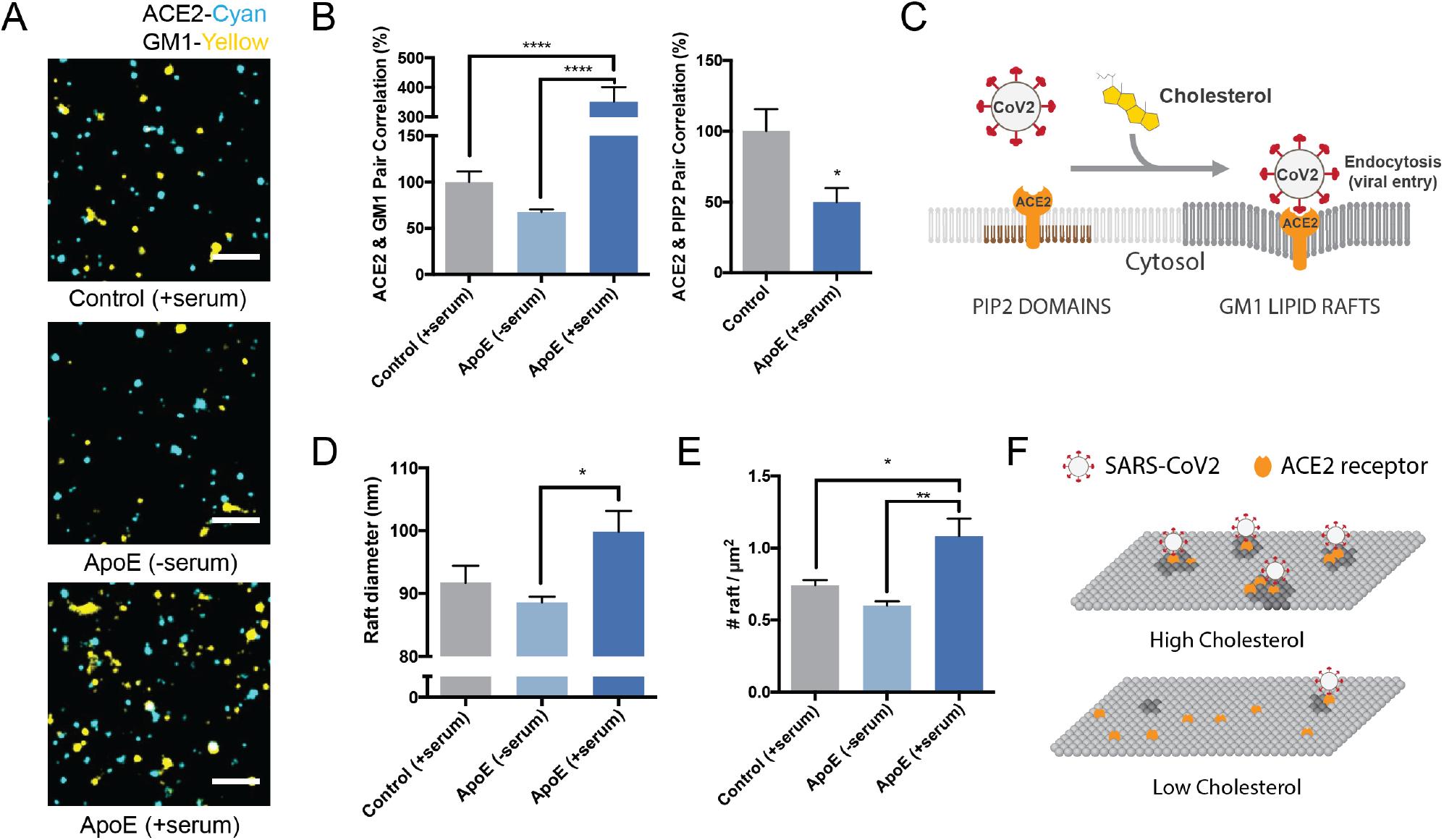
Molecular basis for cholesterol dependent SARS-CoV-2 entry. (**A**) Representative images from dSTORM super resolution imaging. GM1 clusters (yellow) increase dramatically in the presence of blood serum when apoE is present (bottom panel), labelled with CTxB. ACE2 receptor (cyan) labeled with ACE2 antibody has very little overlap with GM1 clusters when blood serum is removed (middle panel). Scale bar, 1 μm. (**B**) Pair correlation from dSTORM imaging. ACE2 was compared to GM1 lipids (CTxB labeled, left) and phosphatidyl inositol 4,5-bisphosphate (PIP2, right). Loading cells with cholesterol (apoE + serum; dark blue shading) increased the pair correlation with GM1 clusters and decreased the pair correlation with PIP2 clusters. Hence ApoE-mediated cholesterol influx moves ACE2 from PIP2 clusters to GM1 clusters. (**C**) Model showing cholesterol-dependent cluster associated protein activation (CAPA) of ACE2 from PIP2 domains (brown lipids) to endocytic GM1 lipids (dark grey) and enhanced SARS-CoV-2 viral entry. (**D**) cluster size analysis suggests GM1 clustering is regulated by apoE mediated cholesterol transportation between serum and cell membrane. (**E**) ApoE transportation of cholesterol regulates apparent cluster density. Cholesterol influx from serum results in an increase in number and apparent size. Data are expressed as mean ± s.e.m., *P<0.05, **P<0.01, ***P<0.001, ****P<0.0001, one-way ANOVA.) (**F**) Model for SARS-CoV-2 viral entry in high and low cholesterol. When cholesterol is high ACE2 translocated to GM1 clusters a position optimal for viral binding and endocytosis. When cholesterol is low, ACE2 traffics away from GM1 clusters and position that poorly facilitates viral infection (PIP2 domains not shown).

Over expression of ACE2 renders HEK293T cells highly susceptible to infection. Given the similar level of ACE2 protein measured in WT HEK293T compared to other endogenous cell types, we investigated the relative effect of ACE2 overexpression in HEK293T cells. We observed ~20 times more ACE2 protein in over expressing cells compared to endogenous systems (Figure 1B) and ~100 times higher SARS-PV infectivity (Figure 1C) compared to WT HEK293T. The non-physiological condition of over-expression combined with the fact that ACE2 levels were the same in VeroE6 and HEK293T cells despite their dramatic difference in infectivity led us to consider the cholesterol-based mechanisms of infectivity.

At a whole membrane level, one function of cholesterol is to cluster membrane proteins^37^ including immune proteins in order to facilitate an immune response^38^. Clustering can also increase viral entry^39,40^ suggesting cholesterol induced clustering could be a factor influencing SARS-CoV-2 entry. To test a potential role for cholesterol we compared highly infectable VeroE6 cells with poorly infected HEK293T cells.

We found that VeroE6 cells had the highest amount of FC, ~20% more than HEK293T (Figure 1B, D). Interestingly, total cholesterol (FC + cholesteryl esters (CE)) was roughly the same (Figure 1E). FC is typically in the plasma membrane whereas esterified cholesterol is more prominent in internal stores^27–29^. We reasoned that cholesterol should produce a change in GM1 lipid clustering in Vero E6 cells v.s. HEK293T cells. To test GM1 clustering, we fluorescently labeled GM1 lipids with cholera toxin B and imaged the average apparent cluster size using direct stochastical optical reconstruction microscopy (dSTORM). dSTORM and other super-resolution techniques are capable of visualizing nanoscale arrangements (i.e. sub-100 nm diameter lipid domain structures) in intact cellular membranes^41–43^. We found Vero E6 have extremely large GM1 clusters, on average 200 nm in diameter, almost twice that of HEK293T cells (Figure 1F). These results suggest that although the total cholesterol level in HEK293T and Vero E6 are the same, the distribution of cholesterol in different cellular compartments are different. Compared with HEK293T, VeroE6 has more cholesterol in the plasma membrane, contributing to GM1 lipid clusters with higher integrity.

### Cholesterol dependent SARS-CoV-2 pseudovirus entry

To directly test the cholesterol dependence of SARS-CoV-2 entry we infected HEK293T cells with retrovirus pseudotyped with the SARS-CoV-2 S protein (SARS2-PV), a segment of the S protein which binds to the ACE2 receptor and recapitulates viral entry^44,45^. Viral particles were prepared fresh to maximize HEK293T infection. Cholesterol was loaded with 4 μg/mL apolipoprotein E (apoE). ApoE is a cholesterol carrier protein linked to Alzheimer’s and the severity of COVID19^46^. In tissue, apoE binds to low-density lipoprotein (LDL) receptor and facilitates loading and unloading of cholesterol into cells (Figure 2A). When apoE is in excess in low cholesterol conditions, it facilitates efflux of cholesterol from the cell^47^ (Supplemental Figure S1B). To provide a source of cholesterol to the apoE, we added 10% fetal bovine serum (FBS), a common source of cholesterol ~310 μg/mL. Importantly, apoE is not present in FBS^48^ allowing us to carefully control cholesterol loading. Cells were treated acutely (1 hr.) with apoE. We adapted the technique from our studies of cholesterol loading in the central nervous system^49^.

We found cholesterol loading (apoE + serum) significantly increased SARS2-PV entry in HEK293T cells. Figure 2B shows viral entry increased 50% compared to control (serum only). Cholesterol efflux (apoE-serum) had the opposite effect; viral entry decreased almost 50% (Figure 2B). Similar effect was also observed in Vero E6 cells (Figure 2C). Methyl-beta-cyclodextrin (M□CD), a chemical known to extract cholesterol from the membrane of cells, inhibited more than 90% of SARS2-PV entry (Figure 2D). To avoid any potential effect of cholesterol directly on the virion, the cells were pretreated for all cholesterol conditions (one hour), washed and then exposed to the virion, i.e., the virion did not experience any cholesterol treatment, only the cells.

VSV-G is an genetically unrelated virus that enter through a cholesterol dependent endocytic pathway. We found infection of cells with a pseudotyped VSV-G had less sensitivity to cholesterol compared to SARS2-PV with identical treatment, although a similar trend was visible as expected for a cholesterol dependent pathway (Supplemental Figure S2A). Likewise, cholesterol modulation with M□CD had a much less effect on VSV-G viral entry compared to SARS-CoV-2 (Supplemental Figure S2B) confirming that SARS-CoV-2 is particularly sensitive to cholesterol.

We also tested the effect of cholesterol on ACE2 activity. Using a fluorescent ACE2 substrate we showed activity increased ~40% with apoE (cholesterol loaded) compared to no apoE (cholesterol depleted), suggesting cholesterol regulates ACE2’s endogenous enzymatic activity is in HEK293T cells.

### Cholesterol dependent trafficking of the ACE2 receptor to GM1 endocytic entry point

In biological membranes we have shown select cluster associated proteins easily move between clusters of GM1 lipids and clusters of phosphatidylinositol 4,5-bisphate (PIP2) lipids with changing cellular cholesterol levels (e.g. phospholipase D2), while other proteins remain clustered with GM1 lipids (e.g. beta and gamma secretase)^31,32,49^. Given the sensitivity of SARS2-PV to cholesterol, we hypothesized that the ACE2 receptor may likewise easily shift between an endocytic pathway and a cell surface mechanism based on cholesterol levels in the plasma membrane.

To determine the amount of ACE2 positioned in a GM1 endocytic pathway, we co-labeled ACE2 and GM1 clusters in HEK293T cells with Cy3b labeled ACE2 antibody and Alexa Fluor 647 conjugated fluorescent CTxB (a GM1 specific toxin), treated cells with apoE/serum, and imaged with two color dSTORM.

Figure 3A shows representative dSTORM images of ACE2 and GM1 clusters from apoE treated cells with and without serum exposure (i.e., cells loaded or not loaded with cholesterol, respectively). The association of ACE2 with GM1 clusters was measured by pairwise correlation. Pairwise correlation measures the probability that two molecules are next to each other. For our experiments we compared the pair correlation at the closest radii (5 nm) as a percentage of control cells without apoE.

**Figure 3.**
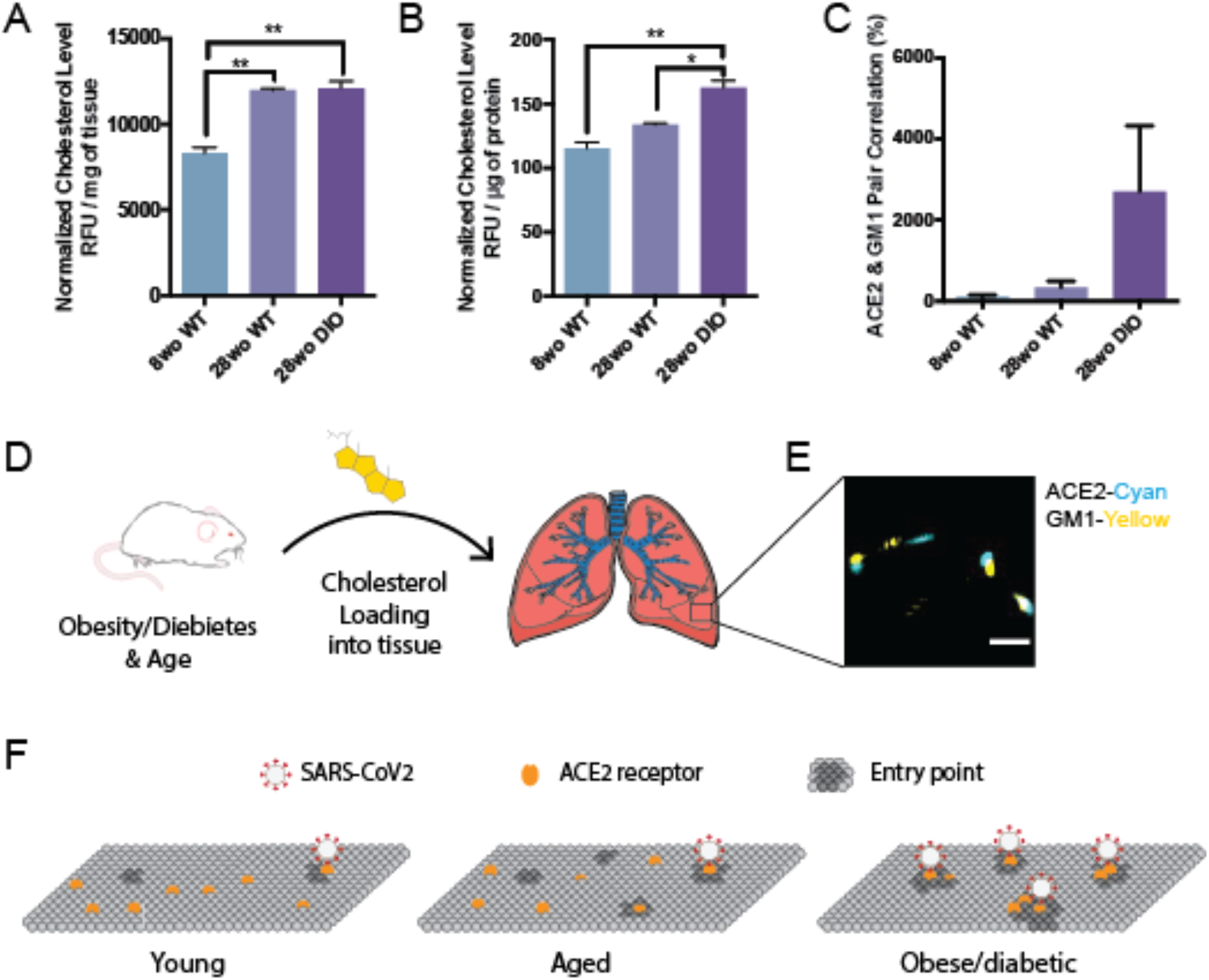
Age and disease dependent cholesterol loading into mouse lung. (**A-B**) Cholesterol measured per mg of lung tissue (A) and per μg of total proteins in the lung (B) in 8-week-old (wo), 28wo mice. An aged diabetic mouse, generated by diet induced obesity (DIO), loaded ~45% more cholesterol into its lung tissue per μg of total proteins. Expressed as mean ± s.e.m., *P<0.05, **P<0.01, one-way ANOVA. (**C**) Association of ACE2 with GM1 clusters determined by dSTORM pair correlation in mouse lung tissue. (**D**) Cholesterol is taken up into the lung tissue of mice fed a high fat diet. (**E**) Representative dSTORM images from (C). Cholesterol (yellow shading) was loaded into the lung of mice (left) and fixed and removed and the tissue directly imaged. Scale bar, 500 nm. (**F**) Proposed model for SARS-CoV-2 infection of lung tissue of obese patients. In young, cholesterol is low, there are few entry points, and ACE2 is ill positioned with PIP2 domains (not shown) away from GM1 clusters (dark grey, entry points). With age, average cholesterol levels in the tissue increases. In obese/diabetic patients, tissue cholesterol levels are the highest and ACE2 receptor is positioned in GM1 clusters for optimal viral entry (PIP2 domains not shown).

We found pairwise correlation of ACE2 with GM1 clustered increased more than 3-fold in cholesterol loaded cells (Figure 3B). We loaded cholesterol using 4 μg/mL apoE and 10% serum, the same as our viral entry assay. The increased correlation was not due to a change in ACE2 concentration on the cell surface (Supplemental Figure S3A-B). As a positive control, and to confirm the effect depends on cholesterol and not another component of the serum or apoE regulation, we treated the cells with M □CD. In agreement with our viral entry assay, M□CD reduced ACE2 localization with GM1 lipids by ~70% (Supplemental Figure S3C).

We then co-labeled ACE2 with an antibody that binds (PIP_2_) lipids. PIP_2_ is a polyunsaturated signaling lipid that forms cholesterol independent domains^50^ in the disordered regions away from GM1 clusters ^32, 50–52^ (See supplemental Figure S1A). As expected, PIP_2_ pair correlation was opposite of GM1, loading cells with cholesterol decreased the association of PIP_2_ and ACE2 (Figure 3B). These data suggest that cholesterol moves ACE2 from PIP_2_ clusters to GM1 clusters (Figure 3C). Can you have gradients of cholesterol within a plasma membrane – current model of high or low cholesterol seems overly simplistic

We next investigated the effect cholesterol has on the number of GM1 clusters in the plasma membrane—presumably an estimate of the number of GM1 labeled endocytic entry points for ACE2. Using cluster analysis, we found cells loaded with cholesterol (+apoE) increased in both the number and apparent diameter of GM1 labeled clusters compared to serum only (not loaded control) (Figure 3D-E). Cholesterol depletion by M□CD also robustly decrease the apparent cluster size (Figure S3D). These decreases caused GM1 clusters to separate from each other measured by Ripley’s H function. (Figure S3E). The cells were fixed prior to labeling to reduce potential artifacts in their diameter size due to CTxB clustering^53^ (see methods).

### Cholesterol loading into lung tissue

For cholesterol to have an effect *in vivo*, the cholesterol must be loaded into the cells where the ACE2 expressing cells are located during an infection. To determine whether there is an age-dependent effect on cellular cholesterol we measured cholesterol levels in young (8-week-old) and aged (28-week-old) lungs from C57BL/6J mice. Lungs were fixed, removed, and 2.5 mg of lung tissue assayed. We found cholesterol in the lung of older adults was elevated (~45%) compared to young mice (Figure 4A). The increase was significant with a P value less than 0.01.

**Figure 4.**
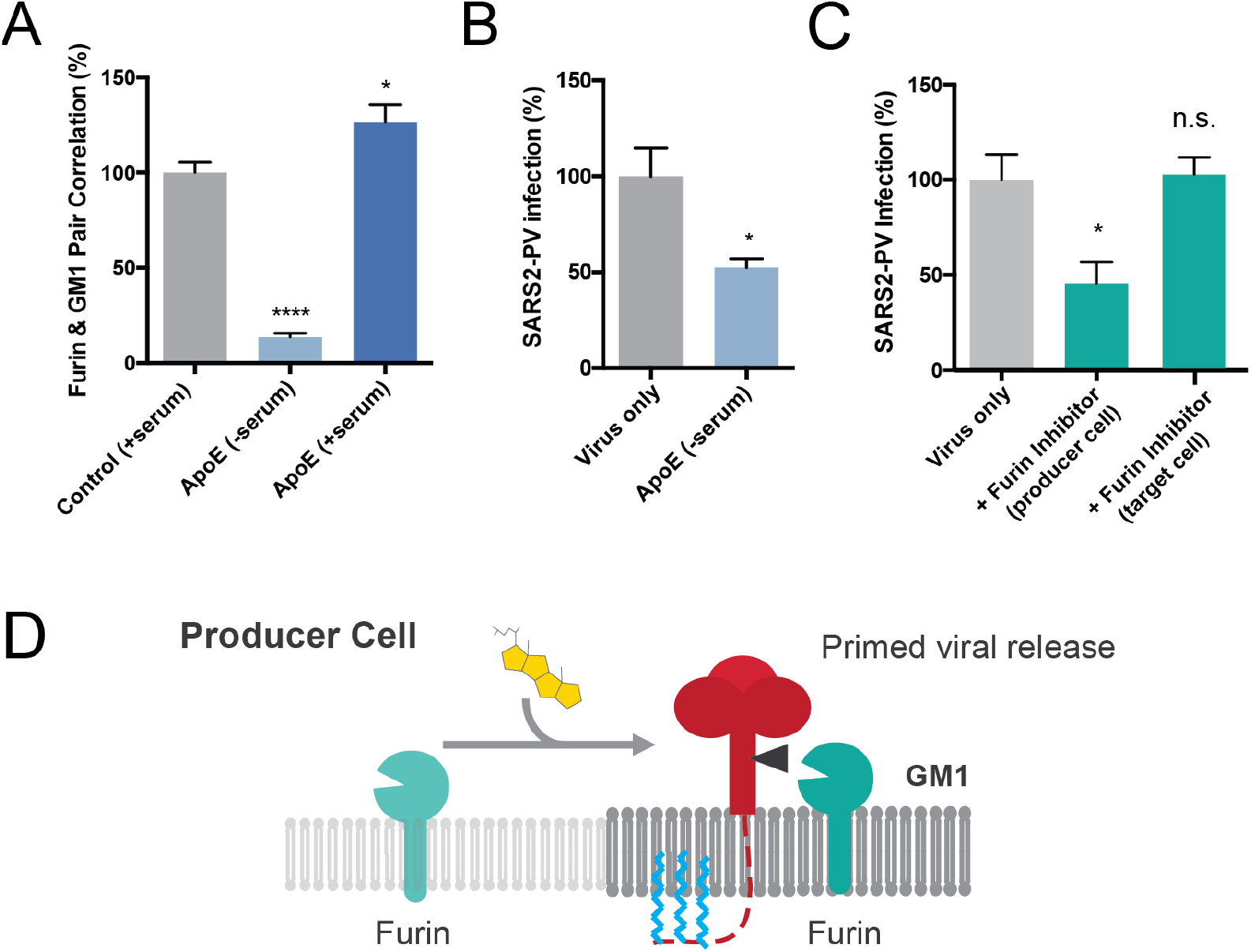
Assembly of viral entrance factors in GM1 lipid rafts. (**A**) Furin localization to GM1 clusters is cholesterol dependent. Furin moves out of the GM1 cluster under low cholesterol condition (apoE extracts cholesterol from membrane). Cholesterol loading by apoE causes an increased furin GM1 lipid pair correlation. Data are expressed as mean ± s.e.m.,*P<0.05, ****P<0.0001, one-way ANOVA. (**B**) Cholesterol depletion with apoE in producer cells decreases the efficiency of viral entry by 50%, suggesting furin’s cluster association is important for spike protein cleavage. Data are expressed as mean ± s.e.m., *P<0.05, two-sided Student’s t-test. (**C**) Furin inhibitor blocks S1/S2 cleavage and virion priming during SARS2-PV exit from producer cells, thus decreasing the efficiency for viral entry. Furin inhibitor treatment in target cells has no effect on viral entry, however when the virus is produced in the presence of a furin inhibitor, infection decreases 50%. The result suggests furin cleavage during virus production increases SARS-CoV-2 viral entry. Data are expressed as mean ± s.e.m., *P<0.05, n.s. P□0.05, one-way ANOVA. (**D**) Speculative role of palmitoylation in SARS-CoV-2 priming of the spike protein. In high cholesterol furin moves from the disordered lipids (light grey lipids) to ordered GM1 lipids (GM1, dark grey lipids). Cholesterol dependent membrane protein translocation is indicated by a grey arrow and the input of cholesterol (yellow shading).

Next we investigated cholesterol levels in lung tissue in aged (28 week) diabetic animals. We increased cholesterol by diet induced obesity (DIO), a well-established mouse model for type II diabetes in of C57B/6J mice ^54^. We noted the weight of the obese lung was increased compared to healthy lung (Supplemental Figure S2C), most likely due to increased fluid permeability in the diabetic mouse. To control for potential excess water in the diabetic mice, we normalized the amount of cholesterol to total protein in the tissue homogenate. Cholesterol normalized per μg of protein was significantly elevated in DIO mice compared to both 8 week and 28-week-old healthy mice (40% and 22% respectively). The increase in lung cholesterol was consistent with the uptake of cholesterol into into cultured cells and suggests an age and obesity induced sensitivity to viral infection exists *in vivo*.

To further confirm a cholesterol dependent effect, we tested ACE2 localization to GM1 clusters in whole lung slices from obese mice. The percent pair correlation for 28wo was more than an order of magnitude higher in lungs from obese mice albeit highly variable (N=10). Nonetheless the trend was consistently higher than either young or control untreated mice (Figure 4C). Labeling in lung was sparser than in our tissue culture (Figure 4E), which likely contributing to the variability. Figure 4F shows an ACE2 based model for age and disease related SARS2 infectivity. In aged obese mice, ACE2 shifts to the GM1 clusters and facilitates viral entry.

### Cholesterol dependent furin localization

During the preparation of pseudovirus, this site appears to be mostly cut in the producing cells^10,18^. For the virus to interact with a protease it must be in proximity (i.e., the same lipid compartment).

Palmitoylation is a post translation modification that attaches a 16 carbon saturated lipid to many proteins and traffics them to GM1 clusters^37^. Palmitoylation of the cysteine-rich endodomain of the SARS-CoV spike glycoprotein is important for spike-mediated cell fusion and viral entry^55,56^. Little is known about palmitoylation of SARS-CoV-2. We aligned SARS-CoV and SARS-CoV-2 and found that all 9 putative palmitoylation sites in SARS-CoV are conserved in SARS-CoV-2 (Figure S4A). A single significant mutation (alanine to cysteine) introduces an additional putative palmitoylation site in SARS-CoV-2, which suggest SARS-CoV-2 remains under evolutionary pressure for palmitate driven viral infectivity.

In a virus producing cell, SARS-CoV-2 palmitoylation is positioned on the intracellular leaflet suitable for targeting the nascent spike protein to GM1 clusters. To test a potential role of cholesterol in facilitating SARS-CoV-2 priming, we loaded producing cells (HEK293T) with cholesterol and imaged furin trafficking in and out of GM1 clusters. Cholesterol (apoE + serum) increased furin’s pair correlation with GM1 clusters, a location that likely favors hydrolysis of the spike protein at the S1/S2 site and facilitates priming of the virus. Unloading cholesterol from the cells (apoE - serum or M□CD) dramatically decreased the percent of furin associated with GM1 clusters (Figure 5A and Supplemental Figure S4B).

A lack of furin priming has been shown to decrease viral entry^18^. If furin priming is decreased with low cholesterol, we expect virion from low cholesterol to also produce less infectious virion. We expressed SARS2-PV in low cholesterol cells pretreated with 4 μg/ml apoE. Figure 5B shows depletion of cholesterol in producer cells yields a much weaker virion. Viral entry into normal control target cells decreased by 50%, suggesting low cholesterol likely decreases priming of the virions.

To confirm reduced cholesterol is similar to inhibited furin in our assay, we produced SARS2-PV in the presence of a furin specific inhibitor and tested viral entry. Entry was reduced ~ 50% for unprimed virus, consistent with previous results and our hypothesis Figure 5C). Adding the inhibitor in the target cell, after the virus is already primed, had no effect (Figure 5C and Supplemental Figure S4C), suggesting furin does not cut an unprimed virion in the target cell.

Figure 5D shows a proposed model for enhanced viral entry in old compared to young due to increased S1/S2 priming in virus producing cells. In young individuals the furin is trafficked outside of GM1 clusters resulting in decreased access of the spike protein to furin. In high cholesterol, furin translocates into GM1 clusters near virus. Low cholesterol in the virion envelope may also contribute to the low infectivity and or recycling of furin into the Golgi (not shown).

### Cholesterol dependent SARS-CoV-2 receptor binding

The effectiveness of viral entry in the disordered region could also result from reduced binding to the ACE2 in the disordered region. To test SARS-CoV-2 binding, we applied the SARS-CoV-2 receptor binding domain (RBD) to HEK293T cells with and without M□CD. M□CD treatment reduced the binding of the RBD (Supplemental Figure S4D). This difference in binding was not seen in SARS-CoV^15^, suggesting the virus may have evolved to bind tighter to GM1 localized ACE2. However, the effect was modest and suggests viral entry rather than binding is likely more affected by cholesterol.

## DISCUSSION

Taken together our findings show the level of cellular cholesterol is a core contributor to SARS-CoV-2 viral entry. A proposed model for age and disease dependent infection of SARS-CoV-2 is shown in Figure 4F. When cholesterol is low (similar to children) there are fewer GM1 entry points, they are smaller, and ACE2 is associated primarily with PIP2 clusters. With age, average cellular cholesterol levels in the lung increases and the number and size of GM1 entry points increases. In chronically ill patients, cellular cholesterol levels increase dramatically and ACE2 receptor shifts to the GM1 entry a location that is presumably more efficient when tissue cholesterol is high (adults + chronic inflammation). The model is supported by data from smokers, age dependent loading of cholesterol into lung tissue of mice (Figure 4A-B), increased size of GM1 clusters in cholesterol and the simultaneous translocation of ACE2 from PIP_2_ domains to GM1 clusters (Figure 3C).

SARS viruses are thought to have two entry mechanisms, surface and endocytic. The nanoscopic trafficking of ACE2 from PIP2 to GM1 clusters suggests the mechanism of viral entry is likely different in young healthy animals compared to chronically inflamed and aged animals. GM1 clusters are clearly part of the endocytic pathway and pertinent during high cholesterol. Whether PIP2 is part of the surface entry mechanism is unknown and would likely be most pertinent to young healthy animals. Over expression of the protease TMPRSS2 greatly facilitates SARS-CoV-2 viral entry^34^. Under normal TMPRESS2 expression, PIP2 may cluster ACE2 away from TMPRSS2. However, the protease does not seem to be necessary, at least not with high cholesterol, as VeroE6 have no detectible TMPRSS2 mRNA and they are easily infected with SARS-CoV-2^34^.

Interestingly, blood cholesterol was found to be lower in the most severe cases of COVID 19 while the cholesterol in the monocytes from the same individuals was elevated^57^. Since we measured free cholesterol in the tissue, we expect a result similar to the blood monocytes, not serum. The high cholesterol in monocytes is consistent with cholesterol loading into tissue during chronic inflammation and suggests blood cholesterol many not always correlate with tissue cholesterol.

ACE2 association with PIP2 lipid domains is very similar to our previous finding with phospholipase D2. PIP2 opposes the function of GM1 clusters by trafficking the proteins away^32,58^. These results suggest that cholesterol is likely in balance with PIP2 in the membrane to regulate the trafficking of an ACE2 in or out of the GM1 cluster and this may change with age and disease.

According to our model, drugs that disrupt GM1 clusters, or drugs that increase cholesterol efflux, are likely to help critically ill COVID19 patients. We and others have shown that polyunsaturated fatty acids, anesthetics, and mechanical force oppose cholesterol signaling through GM1 clusters^31,32,58,59^. Drugs that inhibit cholesterol synthesis, including statins^28^, could also be helpful, especially if taken over time to avoid early infectivity.

## METHODS

### Cell culture

Human embryonic kidney 293T (HEK293T) cells and HEK293T cells overexpressing human ACE2 (hACE2) were grown in Dulbecco’s Modified Eagle Medium (DMEM) containing 10% fetal bovine serum (FBS) and 1% penicillin/streptomycin.

HEK293T cells overexpressing hACE2 were generously provided by Dr. Michael Farzan (Department of Immunology and Microbiology, Scripps Research). Cells were generated by transduction with murine leukemia virus (MLV) pseudotyped with the vesicular stomatitis virus G protein and expressing myc-hACE2-c9, as previously described^56^. Briefly, HEK293T cells were co-transfected by PEI with three plasmids, pMLV-gag-pol, pCAGGS-VSV-G and pQCXIP-myc-hACE2-c9 at a ratio of 3:2:1, and medium was refreshed after overnight incubation of transfection mix. The supernatant with produced virus was harvested 72-hours post transfection and clarified by passing through 0.45μm filter. 293T-hACE2 cells were selected and maintained with medium containing puromycin (Sigma). hACE2 expression was confirmed by SARS2-PV entry assays and by immunofluorescence staining using mouse monoclonal antibody recognizing c-Myc.

### Production of SARS-CoV-2 pseudoviruses

Retroviruses pseudotyped with the SARS-CoV-2 S proteins (SARS2-PV) was produced as previously described^22,23^ with modest modifications as described. HEK293T cells were transfected by X-tremeGENE™ 9 DNA Transfection Reagent (Millipore Sigma, #6365787001) at a ratio of 5:5:1 with a plasmid encoding murine leukemia virus (MLV) gag/pol proteins, a retroviral vector pQCXIX expressing firefly luciferase, and a plasmid expressing the spike protein of SARS-CoV-2 (GenBank YP_009724390) or a control plasmid (VSV-G). All the viral plasmids were provided by Dr. Michael Farzan. Cell culture supernatant containing pseudoviruses was harvested at 72 hours post transfection. For furin inhibitor or apoE treatment, cells were incubated with 25 μM hexa-D-arginine amide (Furin Inhibitor II, Sigma-Aldrich, # SCP0148) or 4 μg/mL apoE (BioVision, #4696-500) respectively, over the process of viral production starting 6 hours after transfection. Virus were then buffer exchanged with Amicon Ultra-4 Centrifugal Unit (Millipore Sigma, #UFC800396) to get rid of furin inhibitor in the media.

### Viral entry assay

HEK293T cells were cultured in 96-well flat culture plates with transparent-bottom (Corning™ Costar™, #3585) pre-coated with poly-D-lysine. Cells were incubated with media containing 4 μg/mL apoE with or without FBS supplementation overnight before viral exposure. For MβCD treatment, cells were incubated with 100 μM MβCD for 30 min prior to virus infection. For furin inhibitor treatment, cells were incubated with 25 μM hexa-D-arginine amide for 1 hour prior to and during the whole process of virus infection.

SARS-CoV-2 pseudoviruses were applied to the cells and allowed to infect at 37 °C for 24 hours. After viral infection, efficiency of viral entry was determined through a firefly luciferase assay. Specifically, cells were washed with PBS once and 16 μL Cell Culture Lysis Reagent (Promega, #E153A) were added into each well. The plate was incubated for 15 min with rocking at room temperature. 8 μL of cell lysate from each well was added into a 384-well plate (Corning™, #3574), followed by the addition of 16 μL of Luciferase Assay Substrate (Promega, #E151A). Luciferase activity measurement was performed on a Spark 20M multimode microplate reader (Tecan). All the infection experiments were performed in a biosafety level-2 (BSL-2) laboratory.

### dSTORM super-resolution imaging

#### Fixed cell preparation

Cells were grown to 60% confluence. Cells were incubated with 4 μg/mL purified apoE protein with or without FBS supplementation (overnight) or 100μM MβCD (30 min) in media. Cells were rinsed with PBS and then fixed with 3%paraformaldehyde and 0.1% glutaraldehyde for 20 min to fix both proteins and lipids. Fixative chemicals were reduced by incubating with 0.1% NaBH_4_ for 7 min with shaking followed by three times 10 min washes with PBS. Cells were permeabilized with 0.2% Triton X-100 for 15 min and then blocked with a standard blocking buffer (10% bovine serum albumin (BSA) / 0.05% Triton in PBS) for 90 min at room temperature. For labelling, cells were incubated with primary antibody (anti-ACE2 antibody (Abcam, #ab189168), anti-Furin antibody (Abcam, #ab3467) or anti-TMPRSS2 antibody [EPR3861] (Abcam, #ab92323)) for 60 min in 5% BSA / 0.05% Triton / PBS at room temperature followed by 5 washes with 1% BSA / 0.05% Triton / PBS for 15 min each. Secondary antibody (donkey anti-rabbit cy3b and CTxB-647) was added in the same buffer as primary for 30 min at room temperature followed by 5 washes as stated above. Cells were then washed with PBS for 5 min. Cell labelling and washing steps were performed while shaking. Labeled cells were then post-fixed with fixing solution, as above, for 10 min without shaking followed by three 5 min washes with PBS and two 3 min washes with deionized distilled water.

#### Lung tissue preparation

Lungs were removed from C57BL/6J wild type and diet induced obesity (DIO) mice. Housing, animal care and experimental procedures were consistent with the Guide for the Care and Use of Laboratory Animals and approved by the Institutional Animal Care and Use Committee of the Scripps Research Institute. Mouse lung tissue slicing and staining were performed as previously described^77^ with minor modifications. Mouse tissues were fixed in 4% paraformaldehyde / 1% glutaraldehyde at 4 °C for 3 days. Tissues were sliced by the histology core. Sections (50 μm) were collected and placed into 24-well plate wells containing PBS. Fixative chemicals were reduced by incubating with 0.1% NaBH4 for 30 min while gently shaking at room temperature followed by three times 10 min washes with PBS. Samples were permeabilized with 0.2% Triton X-100 for 2 hours and then blocked with a standard blocking buffer (10% bovine serum albumin (BSA) / 0.05% Triton in PBS) for 6 hours at room temperature. For labelling, samples were incubated with primary antibody for 3 hours in 5% BSA / 0.05% Triton / PBS at room temperature then 3 days at 4 °C followed by 5 washes with 1% BSA / 0.05% Triton / PBS for 1 hour each. Secondary antibody was added in the same buffer as primary for 3 days at 4 °C followed by 5 washes as stated above. Sample labelling and washing steps were performed while shaking. Labeled lung tissues were then post-fixed with fixing solution, as above, for 1 hour without shaking followed by three 30 min washes with PBS and two 30 min washes with deionized distilled water. Lung slices were mounted onto the 35 mm glass bottom chamber (ibidi, #81158) and 2% agarose were pipetted onto the slice to form a permeable agarose pad and prevent sample movement during imaging.

#### dSTORM imaging

Images were recorded with a Zeiss ELYRA PS.1 microscope using TIRF mode equipped with a pil-immersion 63x objective. Andor iXon 897 EMCCD camera was used along with the Zen 10D software for image acquisition and processing. The TIRF mode in the dSTORM imaging provided low background high-resolution images of the membrane. A total of 10,000 frames with an exposure time of 18 ms were collected for each acquisition. Excitation of the Alexa Fluor 647 dye was achieved using 642 nm lasers and Cy3B was achieved using 561 nm lasers. Cells and brain tissues were imaged in a photo-switching buffer comprising of 1%β-mercaptoethanol (Sigma, #63689), and oxygen scavengers (glucose oxidase (Sigma, #G2133) and catalase (Sigma, #C40)) in 50mM Tris (Affymetrix, #22638100) + 10mM NaCl (Sigma, #S7653) + 10% glucose (Sigma, #G8270) at pH 8.0. Sample drift during acquisition was corrected by an autocorrelative algorithm.

For PIP2 imaging, images were recorded with a Bruker Vutara 352 with a 60X Olympus Silicone objective. Frames with an exposure time of 20 ms were collected for each acquisition. Excitation of the Alexa Fluor 647 dye was achieved using 640 nm lasers and Cy3B was achieved using 561 nm lasers. Laser power was set to provide isolated blinking of individual fluorophores. Cells were imaged in a photo-switching buffer comprising of 1%β-mercaptoethanol (Sigma-Aldrich, #63689), 50 mM cysteamine (Sigma-Aldrich #30070) and oxygen scavengers (glucose oxidase (Sigma, #G2133) and catalase (Sigma, #C40)) in 50mM Tris (Affymetrix, #22638100) + 10mM NaCl (Sigma, #S7653) + 10% glucose (Sigma, #G8270) at pH 8.0. Axial sample drift was corrected during acquisition through the Vutara 352’s vFocus system.

Images were constructed using the default modules in the Zen software. Each detected event was fitted to a 2D Gaussian distribution to determine the center of each point spread function plus the localization precision. The Zen software also has many rendering options including removing localization errors and outliers based on brightness and size of fluorescent signals. Pair correlation and cluster analysis was performed using the Statistical Analysis package in the Vutara SRX software. Pair Correlation analysis is a statistical method used to determine the strength of correlation between two objects by counting the number of points of probe 2 within a certain donut-radius of each point of probe 1. This allows for localization to be determined without overlapping pixels as done in traditional diffraction-limited microscopy. Raft size estimation and raft density were calculated through cluster analysis by measuring the length and density of the clusters comprising of more than 10 particles with a maximum particle distance of 0.1 μm. Ripley’s H(r) analysis was performed to study the distribution of lipid raft clusters.

### Lung cholesterol assay

To measure the relative changes in cholesterol level in lung tissue, we developed an Amplex Red-based cholesterol assay. Lungs were removed from C57BL/6J wild type and diet induced obesity (DIO) mice. Housing, animal care and experimental procedures were consistent with the Guide for the Care and Use of Laboratory Animals and approved by the Institutional Animal Care and Use Committee of the Scripps Research Institute. The left lobe of each lung was homogenized with RIPA lysis buffer. After centrifugation, supernatant was collected for cholesterol and protein measurement. 50 μL of supernatant were mixed with 50 μL of assay solution containing 100 μM Amplex red, 2 U/mL horseradish peroxidase, 4 U/mL cholesterol oxidase and 4 U/mL cholesteryl esterase in PBS in 96-well plates with transparentbottom (Corning™ Costar™, #3585). Relative cholesterol concentration was determined for each sample by measuring fluorescence activity with a fluorescence microplate reader (Tecan Infinite 200 PRO, reading from bottom) with excitation wavelength of 530 nm and an emission wavelength of 585 nm. Subsequently, cholesterol level was normalized by tissue weights and protein concentration. Cholesterol signals were then graphed (Mean ± s.e.m.) and statistically analyzed (one-way ANOVA) with GraphPad Prism software (v6.0f).

### Receptor binding domain (RBD) binding assay

HEK293T cells were seeded into 96-well flat culture plates with transparent-bottom (Corning™ Costar™, #3585) one day before experiment. SARS-CoV-2 RBD with a human Fc fusion was a generous gift from Dr. Michael Farzan (Department of Immunology and Microbiology, Scripps Research). HEK293T cells were incubated with 10 μg/mL RBD peptide in cell culture medium overnight at 37°C in 5% CO_2_. Cells were washed with PBS once before fixation with 3% paraformaldehyde. After blocking with a standard blocking buffer (10% bovine serum albumin (BSA) / 0.05% Triton in PBS) for 90 min at room temperature, RBD binding to the cells were labelled with Alexa 647 goat anti-human IgG (Jackson ImmunoResearch, # 109-606-098) for 6 hours at 4°C. After labelling, cells were washed with PBS twice. Relative RBD binding was determined in quintuplicate for each condition by measuring fluorescent activity with a fluorescence microplate reader (Tecan Infinite 200 PRO) with excitation wavelength of 630 nm and an emission wavelength of 680 nm.

### Statistical analyses

All statistical calculations were performed in GraphPad Prism v6.0. For the Student’s t test, significance was calculated using a two-tailed unpaired parametric test with significance defined as p < 0.05, **P<0.01, ***P<0.001, ****P<0.0001. For the multiple comparison test, significance was calculated using an ordinary one-way ANOVA with Dunnett’s multiple comparisons test.

## Acknowledgements

We thank Michael Farzan for the SARS-CoV-2 spike plasmid, hACE2 overexpression HEK293T cell line, and RBD peptide; Andrew S. Hansen for helpful discussion and reading of the manuscript; Montina Van Meter from histology core and Scott Troutman from the Joseph Kissil Lab (Scripps Research, FL) for support in tissue immunohistochemistry. This work was supported by the National Institutes of Health with an R01 to S.B.H. (R01NS112534) and the US Department of Defense with an Accelerating Innovation in Military Medicine to S.B.H. (W81XWH1810782). We are grateful to the Iris and Junming Le Foundation for funds to purchase a super-resolution microscope, making this study possible.

## Supplemental Figures

**Figure S1.**
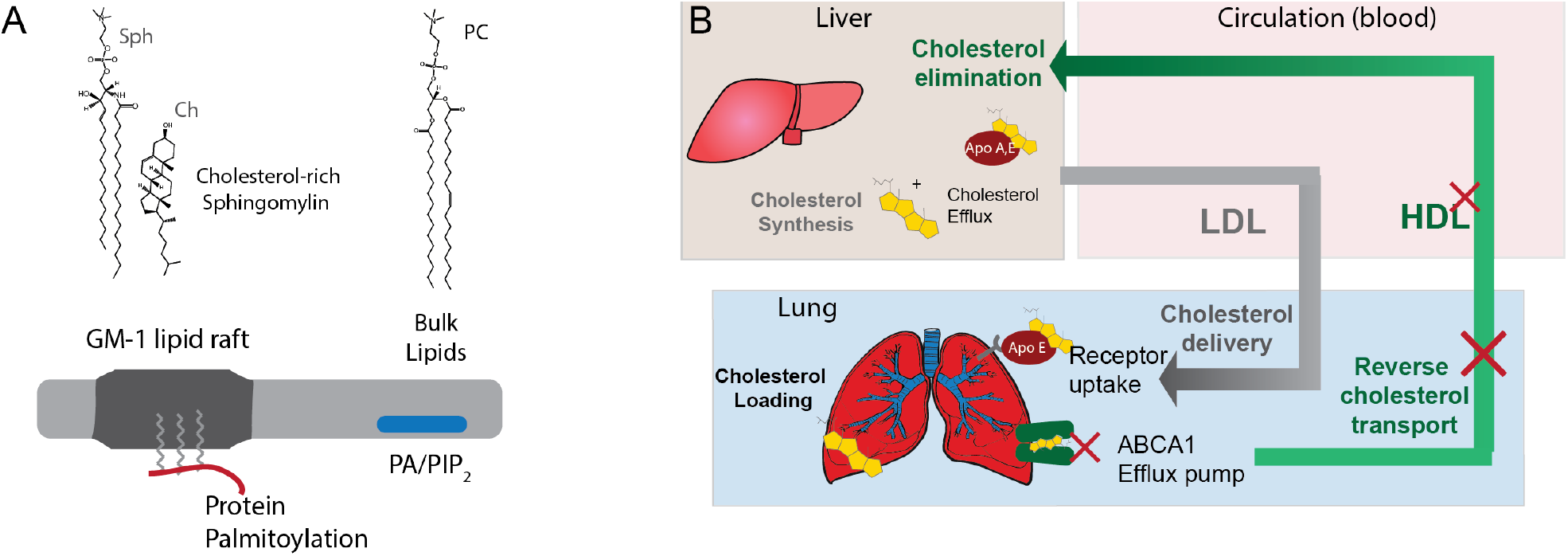
Cholesterol transport and function in GM1 lipid rafts (**A**) The side view of a plasma membrane is shown (top extracellular). The membrane partitions into regions of ordered (saturated) and disordered (unsaturated) lipids. The ordered region contains cholesterol and sphingolipids (Sph). Packing of cholesterol with saturated lipids is thought to provide order and makes ordered lipids thicker than disordered lipids. The disordered region contains unsaturated lipids including phosphatidylcholine (PC), phosphatidic acid (PA), and phosphatidylinositol 4,5-bisphosphate (PIP_2_). PIP_2_ and PA are signaling lipids and PIP_2_ forms its own lipid domains separate from GM1 lipid rafts. Palmitoylation of proteins typically occurs on the cytosolic portion of the membrane and inserts into the inner leaflet (red line with lipids attached). For a virus that exits the cell the palmitoylation may insert into the extracellular leaflet. (**B**) Cartoon of cholesterol loading into lung pneumocyte’s and macrophages. In healthy individuals most cholesterol is produced in the liver. Cholesterol effluxes through ATP binding cassette (ABC) transporters (not shown) and is loaded into apolipoproteins (e.g., A and E; apoA and apoE). The cholesterol is then transported through the blood serum in the form of high- and low-density lipoproteins to the lung. In the lung, receptors located on individual cell membranes (shown as one receptor for simplicity) take up the cholesterol. In healthy individuals, excess cholesterol is effluxed with ABCA1 transporter (green shading) from the plasma membrane of individual cells, loaded into apoA1 or apoE and transported back to the liver for elimination. During chronic inflammation reverse cholesterol transport is inhibited and cholesterol is loaded into macrophage rich tissues in the periphery.

**Figure S2.**
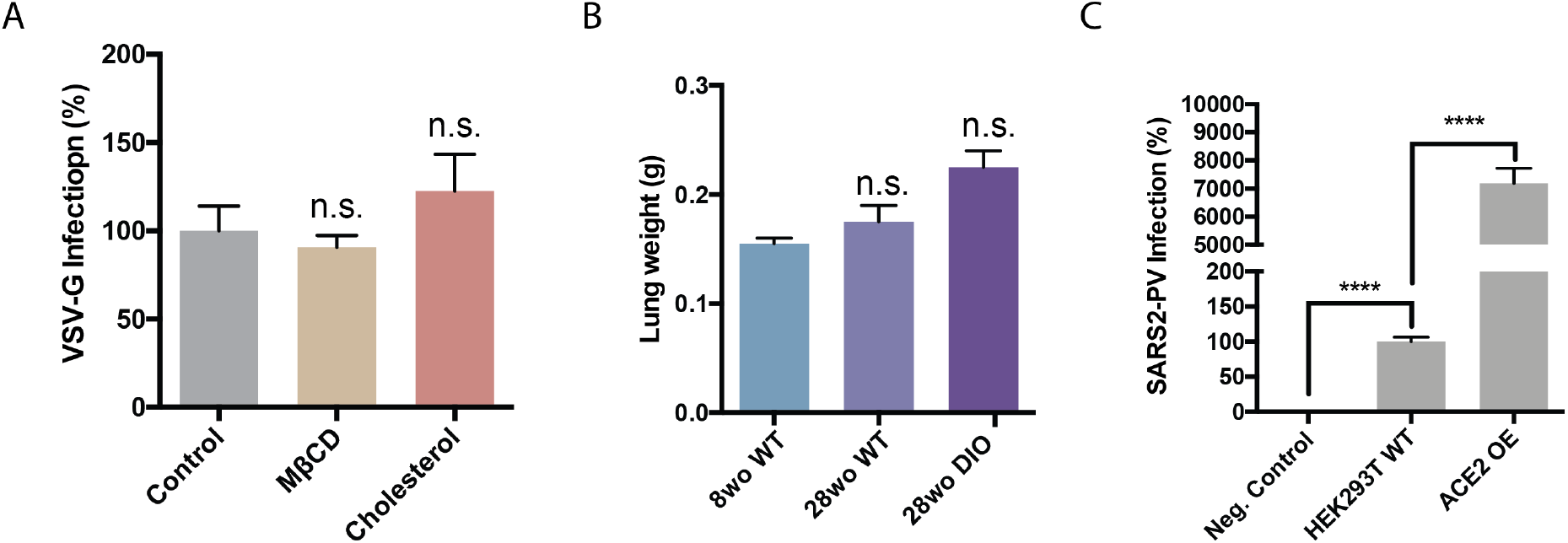
VSV-G viral entry and lung weight. (**A**) Cells were treated with MβCD and a luciferase expressing retrovirus pseudotyped with the VSV-G protein. VSV-G is a similar but distant virus compared to SARS-CoV-2 and serves as a control for age-dependent cholesterol selectivity. Infectivity was monitored by a luciferase activity in cells treated with methyl-beta-cyclodextrin (MβCD) or MβCD with cholesterol. Data are expressed as mean ± s.e.m., n.s.>0.05, one-way ANOVA. (**B**) Lung weight per mg of lung tissue. Lungs were extracted from 8-week-old (wo) and 28wo mice, fixed and assayed for cholesterol with a fluorescent assay. Data are expressed as mean ± s.e.m., n.s.>0.05, one-way ANOVA. (**C**) SARS2 PV viral entry with wild type HEK293T expressing endogenous ACE2 (HEK293T WT) and HEK293T cells overexpressing hACE2 (ACE2 OE). Negative (neg.) control are HEK293T cells treated identical to the other conditions except no virus was applied. The endogenous expression was low but presumably much more physiologically relevant and shows ACE2 is expressed in HEK293T sells albeit at comparatively low quantities. The result is a single side by side comparison, but the values are in a typical range we see for these two systems. Data are expressed as mean ± s.e.m., ****P<0.0001, two-sided Student’s t-test.

**Figure S3.**
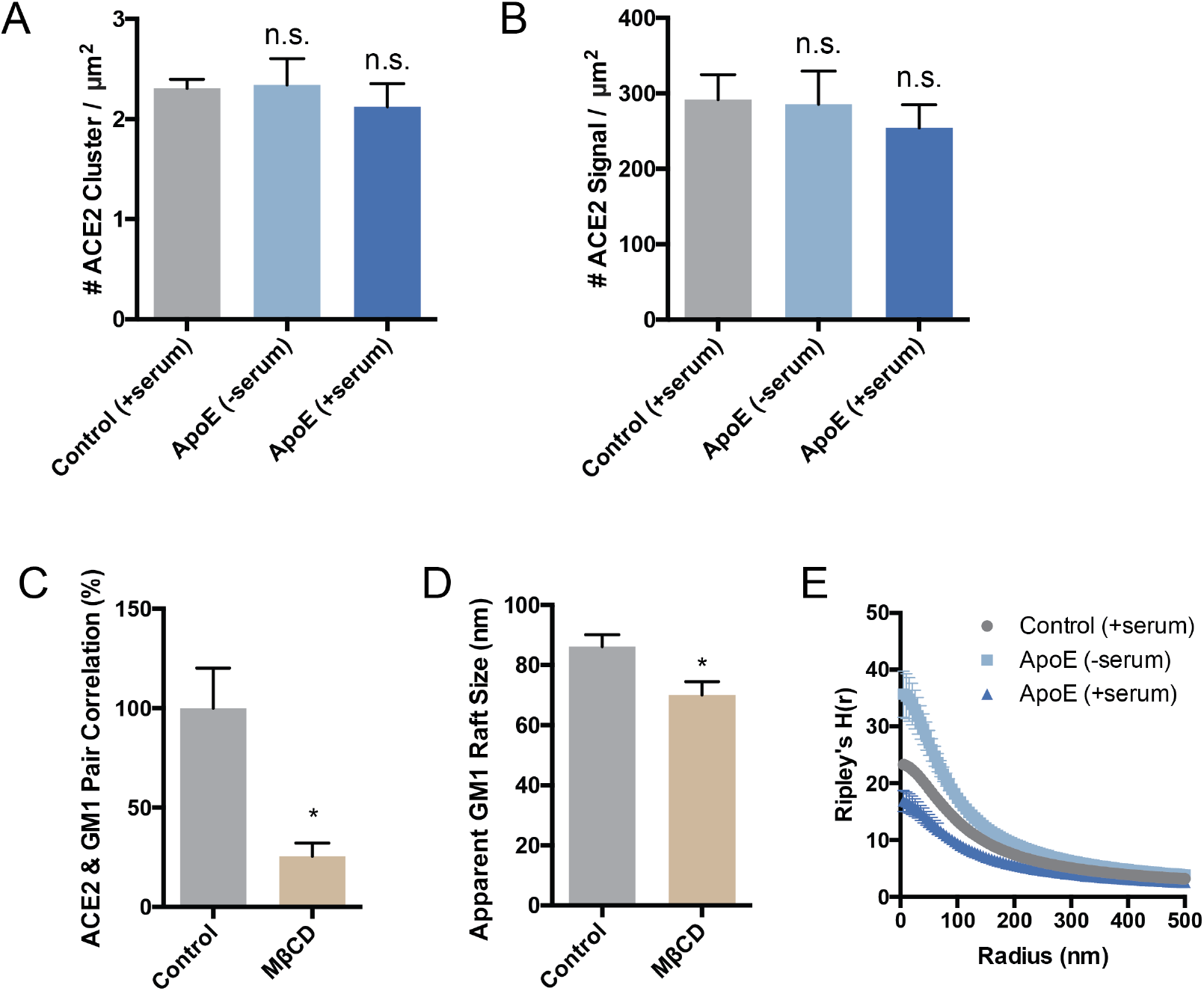
dSTORM of ACE2 cluster (**A**) and signal (**B**) shows that the expression of ACE2 remains roughly constant when cells are loaded or unloaded with cholesterol by the cholesterol transport protein apoE. Although, while not statistically significant ACE2 expression appears to slightly decrease in high cholesterol (ApoE + serum) which agrees with the clinical findings of decreased ACE2 surface expression in older people. Data are expressed as mean ± s.e.m.,*P<0.05, one-way ANOVA. (**C**) Cholesterol depletion with methyl-beta-cyclodextrin (MβCD) diminishes ACE2’s raft localization. (**D**) Cholesterol depletion by MβCD robustly decreases the apparent size of GM1 lipid rafts after CTxB clusterin. Data are expressed as mean ± s.e.m., *P<0.05, twosided Student’s t-test. (**E**) Ripley’s H-Function (H(r)) showing raft separation after cholesterol depletion while distance between rafts gets shorter as apoE transports cholesterol from serum into the membrane.

**Figure S4.**
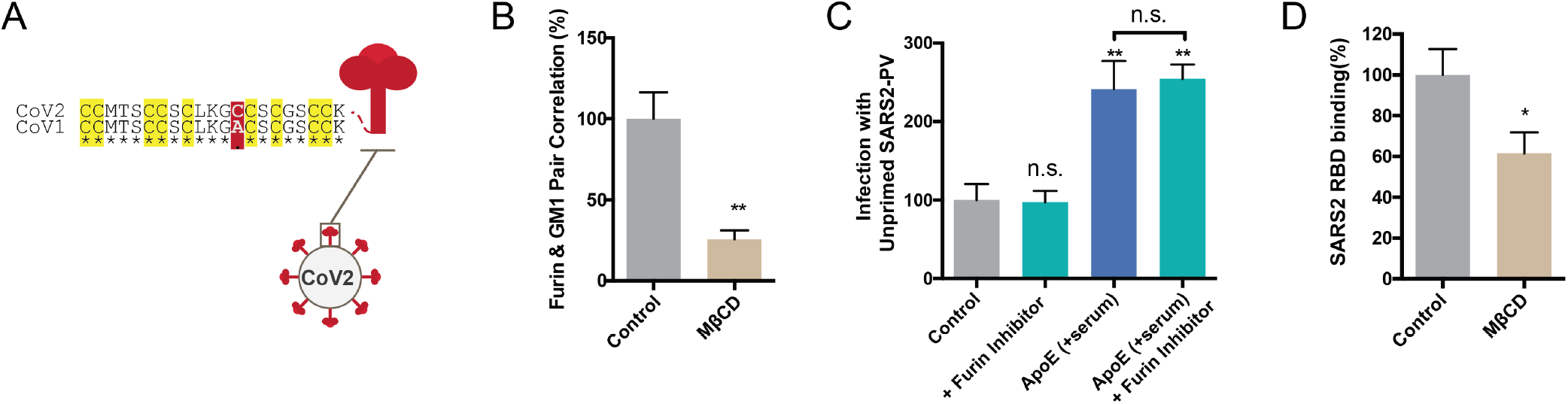
dSTORM of viral processing proteases. (**A**) Sequence alignment of the palmitoylation site in SARS-CoV (GenBank: ABD72995.1) and SARS-CoV-2 (GenBank: QII57161.1). Residues conserved are marked with an asterisk. All 9 putative palmitoylation sites identified in SARS-CoV are conserved in SARS-CoV-2 (yellow highlight). A single mutation introduces a 10^th^ putative palmitoylation site (red shading). (**B**) Cholesterol depletion by MβCD traffics furin out of the GM1 lipid rafts into disordered regions. Data are expressed as mean ± s.e.m., **P<0.01, two-sided Student’s t-test. (**C**) Unprimed virion was produced in HEK cells with furin inhibitor and used to infect cells with and without furin inhibitor and with and without apoE. Furin inhibitor did not change the infectivity of the unprimed virion with or with cholesterol loading (apoE). Hence, furin appears unable to cut the virion in the target cell. Data are expressed as mean ± s.e.m., **P<0.01, n.s. P≥0.05, one-way ANOVA. (**D**) The receptor binding domains (RBD) of SARS-CoV-2 was added to cells treated with or without methyl-beta-cyclodextrin (MβCD). Binding was measured by fluorescence from an Alexa 647 labeled antibody to an Fc tag on the RBD protein.

